# Pathogenic Role of Delta 2 Tubulin in Bortezomib Induced Peripheral Neuropathy

**DOI:** 10.1101/721852

**Authors:** Maria Elena Pero, Cristina Meregalli, Xiaoyi Qu, Atul Kumar, Matthew Shorey, Melissa Rolls, Kurenai Tanji, Thomas H. Brannagan, Paola Alberti, Giulia Fumagalli, Laura Monza, Guido Cavaletti, Francesca Bartolini

**Author notes:** Corresponding author: Francesca Bartolini.

## Abstract

The pathogenesis of chemotherapy induced peripheral neuropathy (CIPN) is still poorly understood. Herein, we found that the CIPN-causing drug, bortezomib (Bort), induces delta 2 tubulin (D2) while affecting MT stability and dynamics in sensory neurons, and that accumulation of D2 is a hallmark of Bort-induced peripheral neuropathy in humans. Furthermore, while induction of D2 was sufficient to cause axonopathy and inhibit mitochondria motility, reducing D2 alleviated both axonal degeneration and loss of mitochondria motility promoted by Bort. Altogether, our data demonstrate that Bort, structurally unrelated to tubulin poisons, can affect the tubulin cytoskeleton in sensory neurons *in vitro, in vivo* and in humans, indicating that the pathogenic mechanisms of seemingly unrelated CIPN drugs may converge on tubulin damage. They further reveal a previously unrecognized pathogenic role for D2 in bortezomib-causing CIPN through its regulation of mitochondria dynamics.

---

CIPN is a “dying back” neuropathy that features a distal to proximal peripheral nerve degeneration widely reported in cancer patients undergoing chemotherapy ^1^. The pathogenesis of CIPN is largely unknown, and this incomplete knowledge is a main reason for the absence of effective neuroprotection strategies to maintain chemotherapy drug anticancer activities while limiting their neurotoxicity ^2-5^. Several classes of anticancer drugs with different antineoplastic mechanisms can induce CIPN ^6-8^. However, sensory impairment ^9^ is the predominant adverse effect associated with each class, suggesting the existence of a common mechanism of pathogenesis ^5^.

Tubulin and microtubules (MTs) are well-established targets for multiple anticancer drugs that can also induce CIPN^10^. The contribution of the MT changes to the onset of CIPN is not well understood, but is strongly implicated as the determining factor. MT hyperstabilization plays a direct role in paclitaxel neurotoxicity ^11^ and a single nucleotide polymorphism in TUBB2a, a gene encoding a tubulin isoform, is directly associated with an enhanced risk of CIPN^12^. Moreover, MT dynamics and stability are markedly influenced by NAD^+^ levels through sirtuin modulation ^13^, suggesting a tubulin-mediated mechanism for the NAD^+^ consuming activity of SARM1 in driving axon degeneration in CIPN models ^14^.

Tubulin, its post-translational modifications (PTMs), MT dynamics and stability play critical roles in neurons, including regulation of long-distance transport, MT severing, Ca^2+^ homeostasis and mitochondria energetics ^14-20^. Each of these functions provide potential mechanisms underlying axon degeneration in CIPN ^21^. Furthermore, tubulin and MTs functionally interact with TRPV1 ^22,23^, and α-tubulin acetylation appears to be a critical component of the mammalian mechano-transduction machinery ^24^. Importantly, anomalies in MT dynamics and tubulin PTMs can drive axon regeneration failure and neurodegeneration ^24-32^.

In addition to taxanes and vinca alkaloids, tubulin changes have been reported downstream of the reversible 26S proteosomal subunit-inhibitor bortezomib (Bort), a widely employed drug with anti-tumor activity in haematological malignancies ^9^. Bort-induced peripheral neuropathy (BIPN) is a painful axonal sensory–predominant and length dependent peripheral neuropathy that affects approximately 40% of the Bort-treated patients through an unknown mechanism ^2^. Interestingly, Bort has been reported to increase MT polymer without disruption of MT ultrastructure, and to affect MT–dependent axonal transport ^9,33-35^. However, a study that analyzes the *in vivo* and *in vitro* effects of Bort on tubulin PTMs and MT behavior as a potential BIPN pathogenic pathway has not been performed. Furthermore, whether perturbation of one or more tubulin PTMs may be sufficient and necessary to induce CIPN has not been determined.

We investigated the effects on tubulin PTMs by acute and chronic administrations of Bort in the cell bodies of dorsal root ganglia (DRG) and sciatic nerves (SNs) isolated from Bort-treated and control rats. In all cases, neuropathy was tested by mechanical and thermal nociception (Fig. S1A and B). Notably, only the chronic regimen appeared to significantly affect both behavior and nerve conduction, and the changes in conduction velocity and evoked response amplitude were more severe in the caudal than in the digital nerve (Fig. S1B). We quantified the fate of selected tubulin PTMs in tissue isolated from chronically treated rats and found that with the exception of polyglutamylated tubulin which appeared to be largely unaffected, all the others were generally reduced, most likely as a result of neurotoxicity induced by chronic dose administrations (Fig. 1C and SFig. 3). When we measured the fate of the same tubulin PTMs in tissues isolated from acutely treated rats prior to any manifested neuronal injury or neuropathic behavior, however, we found that delta 2 tubulin (D2), the only irreversible tubulin PTM and a marker of hyperstable MTs, was significantly increased in SNs and almost 4 fold elevated in DRG cell bodies of Bort treated rats compared to controls (Fig. 1A and B, SFig. 2). Furthermore, when classified by two DRG neuron diameter markers, D2 accumulation appeared higher within small caliber DRG neurons (Fig. 1D and E), phenocopying the cell specificity of the damage to unmyelinated fibers observed in BIPN ^36,37^(Fig. 1D and E).

**Figure 1.**
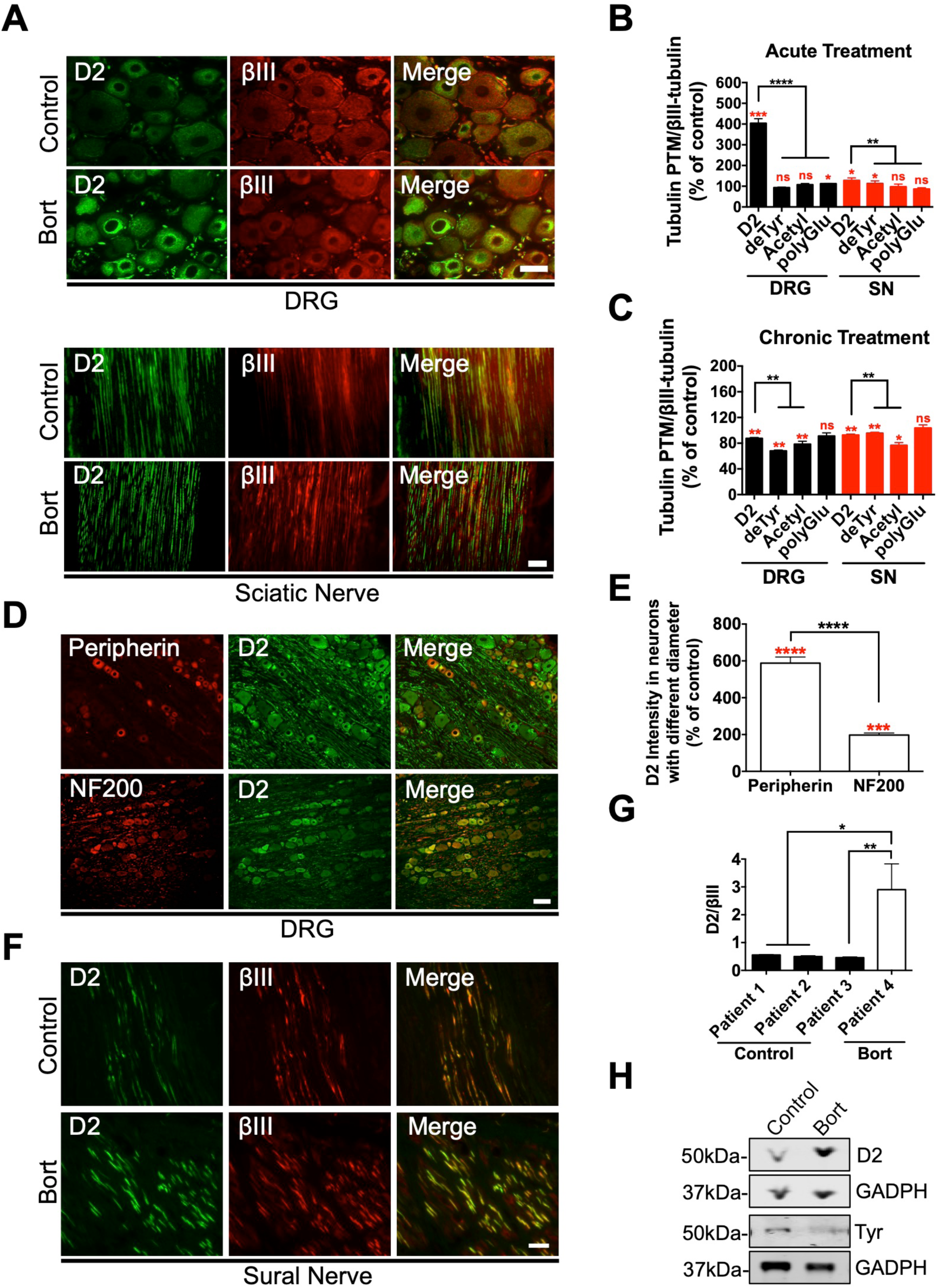
Bortezomib induces D2 levels *in vivo*. **(A)** Representative D2 and βIII-tubulin (βIII) immunofluorescence staining of DRG cell bodies and sciatic nerve (SN) isolated from control rats or rats acutely treated with Bortezomib (Bort) (i.v. 0.2mg/K; 24h). **(B)** Relative tubulin PTM levels measured by quantitative IF in randomly selected DRG cell bodies (n=60-80 per conditions) and sciatic nerve (SN) from rats (n=4 per group) treated with acute (i.v. 0.2mg/K; 24h) or chronic **(C)** (0.2mg/kg, 3x week for 8w) doses of Bort. **(D)** Representative peripherin (smaller unmyelinated C-fibers) and NF200 (large myelinated A-β fibers) immunofluorescence staining of DRG cell bodies from rats treated with Bort for 24h. **(E)** Relative D2 tubulin values measured in DRG neurons positive to either peripherin or NF200. Data are mean values ± SEM, ****,p<0.0001;***,p<0.001;**,p<0.01;*,p<0.05 by ANOVA (B and C in red,) or unpaired two-tailed t-test (B and C in black, E in red) (60-80 DRG neurons per condition). Bar, 20μm. **(F)** Representative D2 and βIII-tubulin (βIII) immunofluorescence staining of sural nerve biopsy from a patient treated with Bort vs control patient. **(G)** Ratio analysis of D2/βIII-tubulin levels measured by IF of fixed tissue from one sural nerve biopsy of BIPN patient vs three sural nerve biopsies from control patients. Data are mean values from 3 sections ± SEM, **, p<0.01 by ANOVA. **(H)** IB of D2 levels in whole cell lysates from one sural nerve biopsy from a patient treated with Bort vs control patient. Tyr, tyrosinated tubulin. GAPDH, loading control.

We tested whether D2 accumulation was promoted by Bort in human disease by examining an available sample from a sural nerve biopsy of a cancer patient affected by BIPN. Indeed, D2 levels were increased in tissue sections and whole tissue lysates of the human sural nerve biopsy from the patient affected by BIPN but not in three control patients, suggesting conservation of the pathways that induce D2 by Bort in humans (Fig. 1F-H).

Next, we investigated whether D2 increase could be detected *in vitro* upon exposure of sensory neurons to pathogenic doses of Bort, and analyzed how D2 levels correlated with axonal injury. To this end, adult DRG neurons grown in culture for 12d, to allow for full recovery from axotomy and for axonal extension, were exposed to Bort up to 72h prior to quantification of relative D2 levels and axonal degeneration from examining individual axons (Fig. 2A-C). D2 was also detected by western blot analysis of whole cell (Fig. 2D and E) and fractionated DRG lysates to evaluate total and partitioned D2 present in soluble and polymerized tubulin fractions (Fig. 2F). Indeed, Bort induced D2 and axonal injury in a dose and time dependent manner with a peak at 48h upon 100nM exposure. A noticeable increase in D2 levels, however, already occurred at 12h prior to the onset of axonal fragmentation at 24h (Fig. 2A-C), and at a dose that did not appear to induce caspase-dependent cellular death up to 48h (SFig. 4). Furthermore, D2 also progressively accumulated in the soluble tubulin fraction starting at 24h (Fig. 2F).

**Figure 2.**
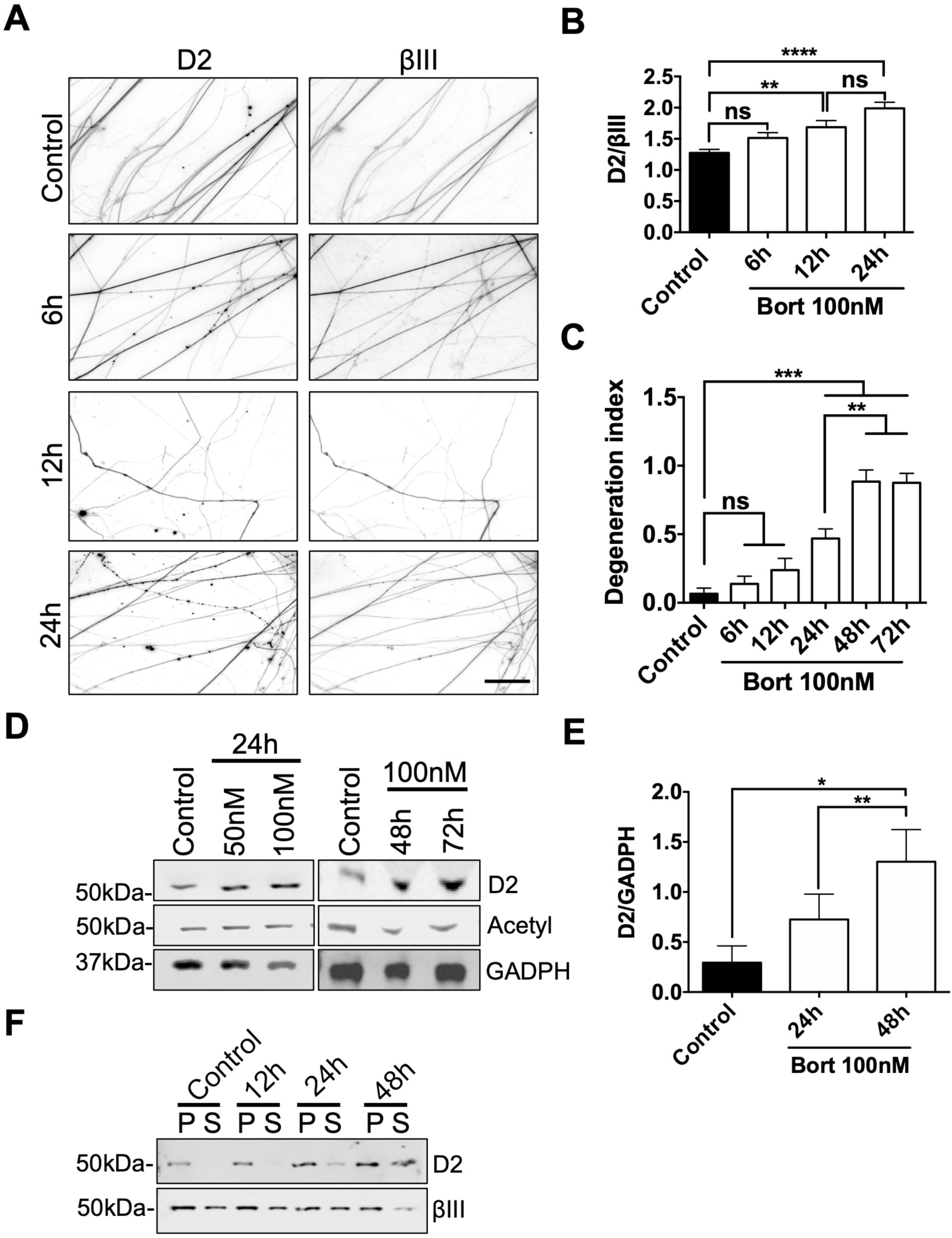
Bortezomib induces D2 at the onset of axonal degeneration. **(A)** Representative IF images of D2 and βIII-tub staining in DRG neurons (12DIV) treated with 100nM of Bort for the indicated times. Bar, 50μm. **(B)** Ratio analysis of D2/βIII-tubulin levels measured by IF of fixed cells treated as in **A. (C)** Time dependent increase of axonopathy in DRG neurons treated with 100nM of Bort for the indicated times. **(D)** IB of D2 levels from whole cell lysates of adult DRG neurons (12DIV) treated with increasing doses of Bort for the indicated times. **(E)** Quantification of normalized D2 levels as in D. **(F)** IB of D2 levels in the MT pellet (P) and soluble tubulin (S) fractions isolated from adult DRG neurons (12DIV) treated with 100nM of Bort for the indicated times. Data are mean values ±SEM from 3 experiments. **, p<0.01, ****, p<0.0001 by t-test (unpaired, two-tailed) (E) and ANOVA (B and C).

To examine the mechanism of D2 induction by Bort, the effects of Bort exposure on the stability of axonal MTs and levels of two rate-limiting enzymes implicated in D2 generation were measured in adult DRG neurons (12DIV) (Fig. 3A-C and SFig.5 A,B). D2 is produced by the sequential action of a tubulin carboxypeptidase that cleaves the last residue of tyrosine from the α-tubulin subunit ^38-41^ followed by the catalytic cleavage of a residue of glutamatic acid by carboxypeptidase 1 (CCP1), a member of a family of tubulin deglutamylases that is expressed in neurons ^42,43^ (Fig. S5A). Tubulin tyrosine ligase (TTL) is a rate-limiting enzyme, which by retyrosinating tubulin controls the abundance of detyrosinated tubulin, a reversible short-lived D2 precursor ^44,45^. *In vitro* Bort acutely induced MT stability measured by the levels of residual β3 tubulin resistant to mild MT depolymerization (Fig. 3A, B) and two tubulin PTMs associated with MT longevity (Fig. 3C). Furthermore, exposure to a Bort dose that resulted in reversible degeneration of peripheral nerve endings significantly promoted an increase in EB3-GFP comet density as seen in live-recordings of nerve fibers in the skin of adult zebrafish (Fig. 3D-F). Endogenous levels of CCP1 and TTL, however, were only slightly affected even at 24h of exposure in cultured DRG neurons, a time point at which significant D2 accumulation and axonal injury begin to be observed (SFig. 5B and Fig. 2).

Altogether, these results demonstrate that in sensory neurons Bort promotes D2 accumulation *in vitro, in vivo* and in humans while acutely affecting MT behavior both *in vitro* and *in vivo*. They further indicate that induction of D2 levels by Bort is the result of acute MT alteration and changes in the longevity of the soluble pool of tubulin caused by proteosome inhibition.

**Figure 3.**
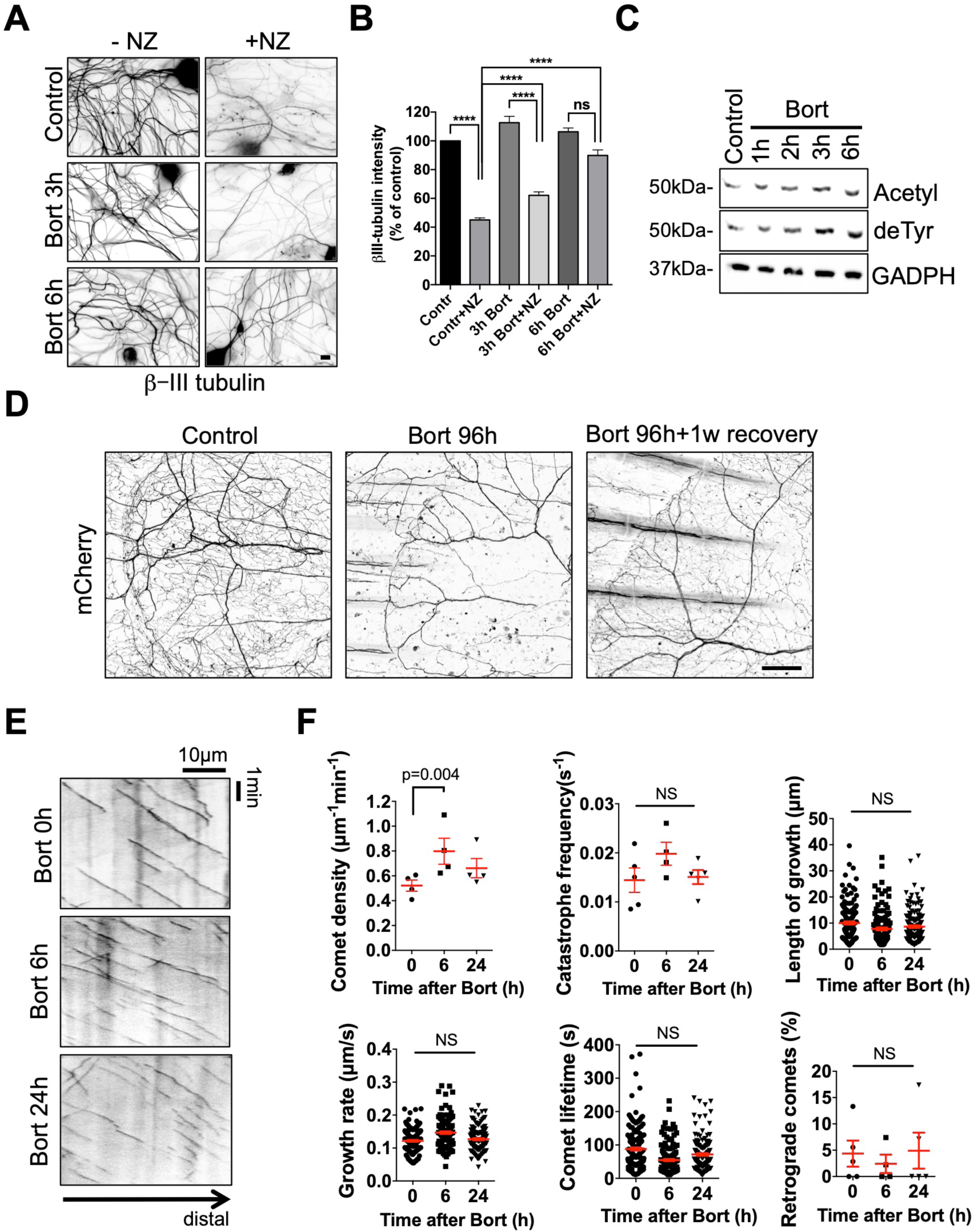
Bortezomib acutely affects MT behavior in DRG neurons *in vit*ro and *in vivo*. **(A)** Representative IF images of βIII-tubulin in adult DRG neurons (12DIV) treated with 100nM of Bort for 3h and 6h and incubated with 1μg/ml nocodazole (NZ) for 45min before MT extraction and fixation. **(B)** Quantification of residual MT mass in proximal axons (100μm) of neurons treated as in A. **(C)** IB of detyrosinated (deTyr) and acetylated (Acetyl) tubulin in cultured adult DRG neurons (12DIV) treated 100nM of Bort for 1-6 h. Data are shown as means ± SEM. ****, P < 0.0001 by t-test (unpaired, two-tailed) and ANOVA. Bar, 10μm. (**D**) Still images from live imaging of mCherry in the peripheral nerve fibers of zebrafish treated with 1.3 uM of Bort for the indicated times. Each image includes endings innervating the surface of a scale. Bar, 100um. **(E)** Representative kymographs at 0h, 6h and and 24h of EB3-GFP comets in the peripheral nerve fibers of zebrafish treated with 1.3uM of Bort. **(F)** EB3-GFP time course analysis of MT dynamics parameters and percentage of retrograde comets in the peripheral nerve fibers of zebrafish treated as in E for the indicated times. Data were pooled from 3 individual fish containing 4 to 5 neurites with up to 80-220 comets and analyzed by Mann-Whitney test. Data are shown as means ± SEM. *, P < 0.05.

To test whether D2 accumulation could alone induce axonopathy, lentiviral delivery of 2 independent TTL shRNA sequences was carried out in untreated adult DRG neurons starting at 5DIV to avoid confounding consequences of TTL loss on axon regeneration ^27^ (Fig. 4A,B). Silencing of TTL was sufficient to increase both D2 and axonal fragmentation in cultured DRG neurons compared to controls (Fig. 4A, B and SFig. 5C). Accordingly, overexpression of recombinant D2 alone but not WT α-tubulin dramatically promoted axonal injury in naive DRG neurons, demonstrating a pathogenic role for accumulation of this irreversible tubulin PTM in the onset of BIPN (Fig. 4C, D and SFig. 5D, E).

**Figure 4.**
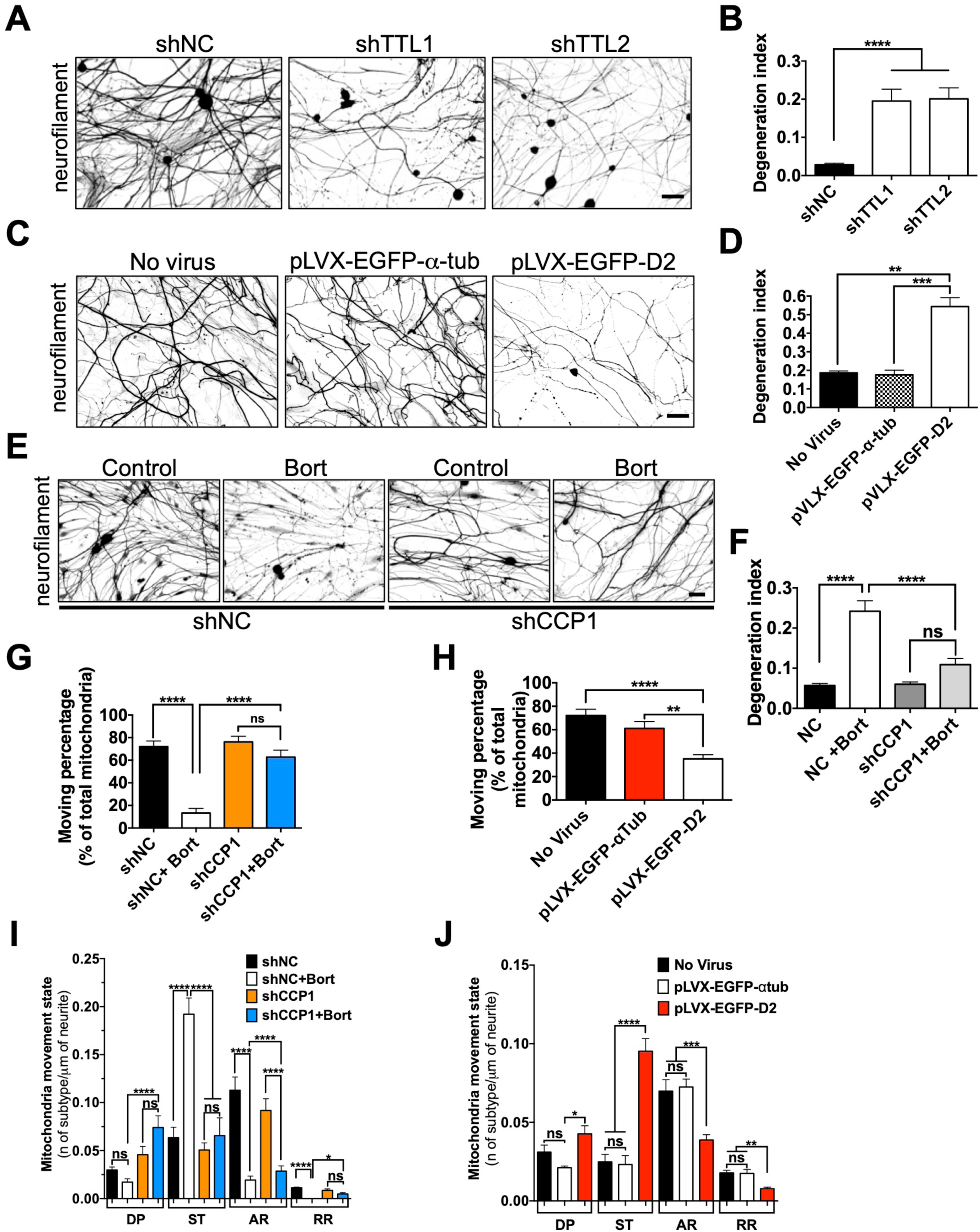
D2 accumulation is sufficient and necessary to drive Bort-induced axonopathy and affects mitochondria motility. (**A**) Representative IF images of DRG neurons (12DIV) silenced of TTL expression for 5 days by lentiviral shRNA delivery. (**B**) Axonal degeneration in DRG neurons treated as in A. (**C**) Representative IF images of DRG neurons (12DIV) transduced with lentiviruses expressing D2-(pLVX-EGFP-D2) or WT-tubulin (pLVX-EGFP-ααtub) (5DIV). **(D)** Axonal degeneration in DRG neurons treated as in C. **(E)** Representative IF images of DRG neurons silenced of CCP1 expression at 2DIV and treated at 10DIV with 100nM of Bort for 24h. **(F)** Axonal degeneration in DRG neurons treated as in E. (**G**) Quantification of the percentage of moving mitochondria in DRG neurons (5DIV) infected at 1DIV with shCCP1 or shNC lentivirus prior 72h infection with mito-DsRed lentivirus followed by Bort treatment (100nM for 24h). **(H)** Quantification of the percentage of moving mitochondria in DRG neurons (5DIV) infected at 1DIV with pLVX-EGFP-D2- or WT-tubulin lentivirus prior to 72h infection with mito-DsRed lentivirus. **(I)** Percentage of different mitochondria movement states, Dynamic Pause (DP), Stationary (STA), Anterograde Running (AR), Retrograde Running (RR) in DRG neurons treated as in E,F. **(J)** Percentage of different mitochondria movement states in DRG neurons treated as in C,D. Data are mean values ± SEM from at least 3 experiments. ***, p<0.001; **, p<0.01, *p<0.05 by ANOVA and unpaired, two-tailed t-test. Bar, 50um.

Next, we examined whether accumulation of D2 was necessary for Bort-induced axonopathy. Lentiviral mediated depletion of CCP1 in differentiated adult DRG neurons promoted substantial loss of D2 with only negligible effects on polyglutamylated tubulin levels, a tubulin PTM recently implicated in neurodegeneration that is negatively regulated by the tubulin deglutamylating activity of CCP1 ^29,46^(Fig. S5F). Under these timely controlled silencing conditions, we found no effect of loss of CCP1 on axonal integrity. Strikingly, however, we observed a significant resistance to axonal degeneration in DRG neurons silenced for CCP1 expression and treated with Bort, indicating a pathogenic role for Bort-induced D2 in driving axonal injury (Fig. 4E, F).

We explored whether the rescue observed in CCP1-depleted cells was intrinsic to the ability of D2 to interfere with mitochondria motility, a hallmark of Bort induced neurotoxicity in DRG neurons ^35,47-49^ and a common feature of axonal injury. Acute depletion of CCP1 to an extent that did not substantially interfere with either D2 or polyglutamylated tubulin levels (SFig. 6A), prevented general loss of mitochondria motility in DRG neurons treated with Bort (Fig. 4G and SFig. 6B), with limited selectivity for dynamic pausing compared to anterogradely or retrogradely running mitochondria pools (Fig. 4I). Accordingly, while WT α-tubulin overexpression did not significantly interfere with any class of motile mitochondria, overexpression of D2 alone, inhibited both dynamic pausing and fast running mitochondria motility in either direction (Fig. 4H, J and SFig. 6C).

In summary, our discovery of the neurotoxic function of accumulated D2 in BIPN provides a new paradigm for the role of deregulated tubulin longevity and modification in the onset of peripheral neuropathy and possibly other related neuropathic conditions. Further work is necessary to detail the mechanisms of this regulation and the cross-talk between D2 and other tubulin PTMs such as polyglutamylation and acetylation in the control of mitochondria energetics and dynamics. It will further be crucial to investigate whether D2 accumulation affects responses to pro-survival and pro-death signals that may play a role in the axon degeneration observed in BIPN, as already suggested in the case of paclitaxel and vincristine ^14,50-52^. Finally, our data provide a strong foundation for testing the potential of targeting tubulin modifying enzymes in drug therapies aimed at preventing and/or rescuing axonal injury observed in CIPN.

## MATERIALS AND METHODS

### Animals

For the *in vivo* experiments, female Wistar rats (Envigo, 175-200g, Udine, Italy) were housed in standard laboratory conditions with *at libitum* access to food and water. Procedures and experimental protocols were compliant with the ethics guidelines described in national (DL.vo 26/2014, Gazzetta Ufficiale della Repubblica Italiana) and international laws (European Union Directive 2010/63/EU: Guide for the Care and Use of Laboratory Animals, U.S National Research Council, 1996). All experiments were approved by the Italian State authorities (# 14/2016). For the *in vitro* experiments, all protocols and procedures used in this study to prepare primary culture of DRG neurons dissected from wild type mouse were approved by the Committee on the Ethics of Animal Experiments of Columbia University and according to Guide for the Care and Use of Laboratory Animals distributed by the National Institutes of Health.

### *In vivo* treatments

Experiments were carried out on female Wistar rats (Envigo, Bresso, Italy), weighing between 175 and 200g at the beginning of the experimental procedure (n=8 rats/group). The rats were housed under constant temperature and lighting conditions with a 12h light/dark cycle and they received food and water *ad libitum*. All procedure was carried out according to the institutional guidelines for animal care in compliance with national (D.L. vo 26/2014, permit No 1161/2016) and international policies (European Communities Council Directive 86/609, Dec 12, 1987; Guide for the Care and the Use of Laboratory Animals, U.S. National Research Council, 1996). Bortezomib (Bort) (LC Laboratories, Woburn, MA) was formulated in 10% Tween 80, 10% ethanol, and 80% sterile saline. The study was divided into two experimental settings. *Setting 1*-acute treatment to test the modifications induced by a single intravenous (i.v) administration of Bort 0.20 mg/kg given as a single dose (and then sacrificed after 24h). *Setting 2*-chronic treatment to evaluate the neuropathic effect induced by Bort injected 0.20mg/kg i.v. for three times/w for 8w as previously described ^53^. For each setting we analyzed untreated animals (Contr) and Bort-treated groups. The electrophysiological and behavioral tests were performed before the beginning of the treatment period (baseline value), 24h after single dose of Bort injection (setting 1), at the end of 8w of Bort treatment (setting 2). All experiments were carried out between 09.00 AM and 04.00 PM.

### Behavioral analysis and electrophysiology

At baseline, 24h after a single Bort administration and 24h after completion of the antineoplastic drug, the animals were tested by a blinded examiner and in random fashion with behavioral methods to evaluate their mechanical and thermal nociceptive thresholds through Dynamic and Plantar tests, respectively, as previously described^54^. Mechanical allodynia in the rat was determined with a Dynamic analgesimeter (Ugo Basile, Comerio-Varese, Italy) which generates a linearly increasing mechanical force. Briefly, a metal filament applied a linear increasing pressure under the plantar surface of the hind limb using an electronic Von Frey hair unit. The withdrawal threshold was measured by applying force ranging from 0 to 50g within 20s. Mechanical threshold reflex was measured automatically by the grams until withdrawal in response to the applied force. Two h after dynamic test, change in thermal hyperalgesia after single and chronical treatments of Bort were monitored using Plantar test (Hargreaves’ method; Ugo Basile, Comerio-Varese, Italy), as described previously ^55^. In brief, after habituation, a movable infrared heat source (IR 50mV/m^2^) was located directly under the plantar surface on the hind paw, and withdrawal latency was used for data analysis. Sensory conduction studies (sensory nerve action potential and conduction velocity) of caudal and hind limb digital nerves (extensively used in the classification of peripheral neuropathies) were assessed orthodromically ^56^ through the Myto2 EGM device (EBN Neuro, Firenze, Italy). Briefly, two sensory nerves (caudal and digital nerves) were assessed by placing recording cathode and anode at 6 and 5cm from the tip of the tail, the ground electrode at 2.5cm from it, and stimulating anode and cathode respectively at 2 and 1cm to measure the caudal nerve alteration. To study digital nerve, recording cathode and anode were placed at the base and at the tip of the fourth toe of the hind-limb respectively, the ground electrode subcutaneously in the sole, and stimulating anode and cathode at the ankle and subcutaneously near the patellar bone, respectively. Intensity, duration and frequency of stimulation were set up in order to obtain supramaximal results. Animals were anesthetized with 2% isofluorane and the body temperature was maintained at 37 ± 0.5°C with a temperature-controlled heating pad (Harvard Apparatus, Holliston, US).

### Quantitative immunoblotting and immunocytochemistry of tubulin PTMs and MT markers

Whole cell lysates cultured DRG neurons were obtained by manual homogenization in SDS-loading buffer followed by sonication (Bransor-Sonifier 250 for 15s), boiling (95°C for 5min) and pre-clearing (5min at 10,000xg) prior to loading and transfer. Protein bound nitrocellulose membranes were probed for detyrosinated (rabbit, 1:200 IF, 1:2000 WB, Millipore), tyrosinated (rat, 1:400 IF, 1:2000 WB, Millipore), acetylated (mouse, 1:300 IF, 1:2000 WB, Sigma Aldrich), D2 (rabbit, 1:400 IF,1:1000 WB, Millipore), poly-glutamylated (1:2000 PolyE; PolyGlu Modification mAb GT335; Adipogen), βIII (mouse and rabbit, 1:500 IF; 1:2000 WB, Abcam), DM1A tubulin (mouse 1:2000 WB, Sigma Aldrich), CCP1 (rabbit, 1:1000 WB, Proteintech), TTL (1:1000 WB, Proteintech), GADPH (mouse and rabbit, 1:5000 WB, TermoFisher and Sigma Aldrich), pheripherin (mouse, 1:200 IF, Abcam), and NF-200 (chicken and mouse, 1:200 IF, Aves Labs) levels using commercially available specific primary antibodies. For WB analysis, secondary antibodies were conjugated to IR680 or IR800 (Rockland Immunochemicals) for multiple infrared detection. Image acquisition was performed with an Odyssey imaging system (LI-COR Biosciences, NE) and analyzed with Odyssey software. For immunofluorescence (IF) studies of fixed tissue specimens, DRG and sciatic nerve sections were processed as follows: paraffin embedded blocks of DRG (L4-L5) and sciatic nerve samples were serially cut (5μm) and deparaffinized by immersing the slices in xylene (7min x 2) followed by 100%, 90%, 80%, 60%, 40% EtOH (5min each) and H_2_O (5min) immersion. Slices were then rinsed with PBS1X (10min x 2) and antigen retrieval carried out by boiling the slices in citric acid solution (10mM, pH 6) for 15min. Sections were cooled down for 15min, rinsed 3 times with PBS 1X, permeabilized with 0.01% Triton X-100 for 10min and blocked with NGS (Thermofisher) 10% in PBS 1h at R.T. Primary antibodies incubations were performed o/n at 4°C in NGS 10% PBS 1X buffer. Alexa Fluor fluorescent dyes-conjugated secondary antibodies were diluted 1:200 in 10% NGS-PBS1X and added for 1h at R.T. For IF studies of isolated DRG neurons, cells were fixed with 4% PFA, followed by membrane permeabilization with 0.1% Triton X-100 and then staining with specific primary and secondary antibodies as described in^28^. Acquisition was performed using spinning disk confocal (Olympus DSU) coupled to a camera controller Hamatsu (C10600 ORCA-R^2^) or epifluorescence microscopy (Olympus IX81) coupled to a monochrome CCD camera (Sensicam QE; Cooke Corporation) and all images were analyzed by ImageJ software. Ratiometric analysis of tubulin PTM/bulk tubulin levels on DRG cellular bodies, SN and proximal and distal axons were performed on randomly selected cellular bodies or axons from images of fixed and immunostained DRG neurons using ImageJ software.

### Isolation of adult DRG neurons

Dorsal root ganglia (DRG) were dissected from 8 to 10w old C57Bl/6J mice in cold HBSS (Life Technologies) or DMEM (Life Technologies) and dissociated in 1mg/mL Collagenase A (Sigma) for 1h at 37 °C, followed by 0.05% trypsin (Life Technologies) digestion for 5min at 37°C and washing with Neurobasal medium (Invitrogen) supplemented with 2% B-27 (Invitrogen), 0.5mM glutamine (Invitrogen), FBS (Sigma) and 100U/mL penicillin-streptomycin. DRG neurons were then triturated using Glass Pasteur pipettes and neuronal bodies resuspended in Neurobasal medium prior to plating onto 12 well plates (over 18mm coverslips) that had been coated o/n with 100μg/mL poly-D-lysine (Sigma) at 37°C and 10μg/mL laminin (Life Technologies) for 1h at 37 °C. At 2DIV Neurobasal medium without FBS was added to the plate. At 4DIV, 10μM AraC (Sigma) was added to media every 4 days.

### MT fractionation assays

DRG neurons were treated with Bort for 24h. At the end of the incubation time, cells were gently washed with warm PHEM buffer (60mM Pipes, 25mM Hepes, 10mM EGTA, and 4mM MgCl2, pH 6.9) once before extraction with PHEM buffer supplemented with 0.05% Triton X-100, protease inhibitor cocktail, and 10μM taxol. After 30min at 37°C, the supernatant containing the soluble tubulin fraction was quickly collected in 20μl 5X laemmli buffer and 100μl 1× PHEM laemmli buffer was added to each tube to collect the MT pellet ^28^.

### Lentiviral infection of isolated adult DRG neurons

Lentiviral particles delivery technique was employed to infect adult DRG neurons with lentiviral cDNA or shRNA. Production of lentiviral particles was conducted using the second generation packaging system as previously described ^57^. In brief, HEK293T cells were co-transfected with lentiviral cDNA or shRNA constructs and the packaging vectors pLP1, pLP2, and pLP-VSV-G (Thermofisher) using the calcium phosphate transfection method. 24, 36 and 48h after transfection, the viral containing supernatant was collected, and the lentiviral particles were concentrated (800-fold) by ultracentrifugation (100,000 x g at 4°C for 2h) prior to aliquoting and storage at -80°C.

### cDNA and shRNA constructs

Lentiviral constructs to overexpress WT and D2 tubulins were generated by subcloning of WT and D2 tubulins from WT and D2 pEGFP-tubulin vectors (Clontech) into pLVX lentivector. Briefly, pEGFP-D2 tubulin was generated by site-directed mutagenesis kit (Agilent) and confirmed by standard Sanger sequencing (Genewiz). Lentiviral WT and D2 tubulins were then generated by digesting pEGFP WT or D2 tubulin with restriction enzymes Afe1 and BamH1 and ligation of the insert into pLVX lentivector.

Lentiviral MitoDsRed was purchased from Addgene (cat n 44386). Lentiviral EB3-EGFP was generated by subcloning EB3-EGFP (a gift from Dr. Polleux) into pLVX lentivector by Afe1 and Not1 sequential digestions.

Two TTL shRNA sequences were purchased from SIGMA in pLKO.1 vectors: shTTL1 (catalog number TRCN0000191515, sequence: CCGGGCATTCAGAAAGAGTACTCAACTCGAGTTGAGTACTCTTTCTGAATGCTTTTTTG) and shTTL2 (catalog number TRCN0000191227, sequence: CCGGCTCAAAGAACTATGGGAAATACTCGAGTATTTCCCATAGTTCTTTGAGTTTTTTG) ^58^. A CCP1 shRNA was purchased from Sigma in pLKO.1 vector (catalog number TRCN0000347379, sequence: CCGGTGGAGTGAAACAGCTTATTATCTCGAGATAATAAGCTGTTTCACTCCATTTTTG). The pLKO.1 vector with noncoding (NC) sequence (Sigma) was used as control.

### MT stability assays

DRG neurons (12DIV) were treated with Bort for 3h and 6h before addition of 1μg/ml nocodazole for 45min. At the end of the incubation time, DRG neurons were gently washed with warm PHEM buffer (60 mM Pipes, 25mM Hepes, 10mM EGTA, and 4mM MgCl2, pH 6.9) once before extraction with PHEM buffer supplemented with 0.05% Triton X-100, protease inhibitor cocktail, and 10μM taxol. After 5 min at 37°C, a matching volume of 2× fixative buffer (8% PFA and 0.2% glutaraldehyde in 1× PHEM) was added dropwise to the coverslips, and then cells were incubated for another 30 min at 37°C. DRG neurons were finally washed with PBS and processed for IF labeling. All images were analyzed using ImageJ software by measuring the mean intensity of major proximal neurites (within 100 μm from the cell body). Data are mean βIII-tubulin levels expressed as percentages of vehicle control ± SEM.

### MT dynamics assays in zebrafish nerve endings

Adult zebrafish (*Danio rerio*) were raised in an Aquaneering fish habitat set to pH 7.5, 400-500us salinity and 28C with 10% water exchanges daily. Fishes used in the experiment were in a Nacre mutant background and doubly heterozygous for the transgenic insertions P2X3.LexA.LexAop.EB3-GFP and P2X3.LexA.LexAop.mCherry. The mCherry labeled strain was a generous gift from Dr. Alvaro Sagasti of UCLA and the EB3-GFP fish were generated in the Rolls lab using the gateway system and tol2 kit as previously published in ^59^. After the first imaging session fish were housed in standard mating cages, one fish per cage with no divider, and their water exchanged every 24h with fresh system water with 0.01% DMSO and 1.3µM Bort, or 0.01% DMSO for vehicle treated control fish. Fish were mounted in 2% low melt agarose, anesthetized with 0.012% tricaine but otherwise were intubated and recovered as describes by Rasmussen et al. 2018^60^. Imaging was performed on a Zeiss LSM800 inverted microscope using a 20x 0.8NA air objective. Time series of images was taken with a zoom of 5.3, 512×512 pixel size and a 4us pixel dwell time for a frame time of 1.3s. Kymographs were generated using the multi kymograph feature of ImageJ. All zebrafish experiments were performed after receiving approval from the Penn State Institutional Animal Care and Use Committee, and guidelines on euthanasia and pain management were followed.

### Neurodegeneration index measurements in DRG neurons

Images of six random fields of dissociated adult DRG neurons (12DIV) fixed and immunostained with mouse anti-neurofilament (2H3-s) antibody (1:300, DSHB) were acquired using a 20X objective lens (Olympus IX81) coupled to a monochrome CCD camera (Sensicam QE; Cooke Corporation). To quantify axonal degeneration, the area occupied by the axons (total axonal area) and degenerating axons (fragmented axonal area) was measured in the same field from images of DRG neurons treated with the drug up to 72h. Images were binarized and fragmented axonal area measured by using the particle analyzer module of Image J (size of small fragments = 20–10,000 pixels). Degeneration index was calculated as the ratio between fragmented axonal area and total axonal area ^61^.

### Mitochondria motility assays

Dissociated adult DRG neurons (2DIV) were transduced with shCCP1, shNC or pLVX-D2 and pLVX-α-tubulin lentiviruses; 24h later neurons were transduced with Mito-DsRed lentivirus and imaged at 5DIV using an epifluorescence microscope (Nikon Ti) equipped with a controlled temperature and CO2 incubator. The pLKO.1 vector with noncoding (NC) sequence was used as control for shCCP1 while non-infected neurons were used as control for pLVX-D2 and pLVX-α-tubulin lentiviruses. Proximal axons (within 100μm from the cell body) were selected for imaging and movies were acquired at 10s/f for 30min. Mitochondria motility was measured using kymograph analysis in ImageJ. Mitochondria were further classified into 2 different movement states, stationary (ST) and dynamic. Mitochondria were defined ST if remained in an immobile state for the entire length of the movie. Dynamic mitochondria were subclassified as dynamic pausing (DP) if showing persistent and short oscillatory movements (0.3-5um), anterograde (AR) and retrograde (RR) running if moving for short or longer (>5um) distances in either direction. Number of different mitochondria movement states per movie was manually scored and quantified on kymographs.

### Caspase Cleavage Assay

Caspase activation was assessed in DRG neurons treated with 100nM of Bortezomib for 12, 24 and 48h using an *in vitro* caspase-3 like cleavage assay utilizing the chemical substrate DEVD-7-amino-4-methylcoumarin (AMC) (Enzo Life Sciences, Framingdale, NY #ALX-260-031). At the end of treatment, neurons were lysed prior to the addition of AMC substrate and incubated for 30min at 37°C in the dark. The generation of the fluorescent AMC cleavage product was measured at 360nm excitation and 460nm emissions, using a fluorescence plate reader (SpectraMax i3x multi-mode detection platform, Molecular Devices). Staurosporine addition at the concentration of 600nM for 24h was used as a positive control ^62^.

### Sural nerve biopsies and clinical history of control and BIPN patients

Clinical history of the BIPN patient shown was previously described ^63^. Briefly, the patient was administered bortezomib 1.3gm/m^2^ and a second dose after 4 days to treat a marginal zone lymphoma. Neurological examination showed distal weakness, which was more severe in the legs, and sensory loss to pin, temperature and vibration distal in the hands and feet. An EMG and nerve conduction study showed signs of a sensorimotor neuropathy, with axonal loss and multifocal demyelination. The patient’s sural nerve biopsy showed a significant axonal loss involving both large and small myelinated fibers, with no robust axonal regeneration. There were scattered histiocytes/macrophages associated with active axonal breakdown in the nerve fiascicles, along with perivascular T-cell-dominant lymphocytic inflammation. No overt vasculitis or amyloid deposition was seen. By teasing fiber analysis, 22% of the teased myelinated fibers showed Wallerian degeneration, 22% myelin changes in the forms of segmental demyelination and remyelination. After stopping bortezomib nerve conduction studies after 22 months showed improvement in the compound muscle action potential amplitudes. One control patient had a right sural nerve biopsy for a suspected mitochondrial disease (not specified). The morphology revealed a peripheral nerve with mild changes without significant inflammation, consisting of minimal axonal loss with 10% of the teased fibers exhibiting segmental remyelination. No amyloid deposition was observed. The second control patient was a suspected case of idiopathic peripheral neuropathy, not due to neurotoxic medications. The nerve biopsy, however, showed minimal changes that were indistinguishable from age-related alteration with minimal axonal loss and 9.3% of the teased fibers exhibiting segmental remyelination. No vasculitis, amyloidosis, and immunoglobulin deposition were detectable. The third control patient underwent a sural nerve biopsy with the symptoms of peripheral neuropathy, parkinsonism, bulbar palsy and history of Lyme disease. The sural nerve revealed minimal changes, without any significant axonal loss. By the nerve teasing, 11% of the fibers show segmental remyelination. No vasculitis or amyloid deposition was identified. The sural nerve biopsies were performed as previously described ^64^.

### Statistics

Statistical analyses of immunolabeling of tubulin PTM/bulk tubulin levels (delta 2, detyrosinated, acetylated, polyglutamylated, βIII, total α), immunoblotting and neurodegeneration index were conducted by two-tailed unpaired t-test (parametric and non parametric) or ANOVA for comparison of multiple samples. MT dynamics in zebrafish were analyzed by Mann-Whitney test. Each set of experiments was performed at least 3 times. Data were reported as mean ± SEM. Statistical significance was set at p<0.05.

## SUPPLEMENTAL MATERIAL

**SFigure 1.**
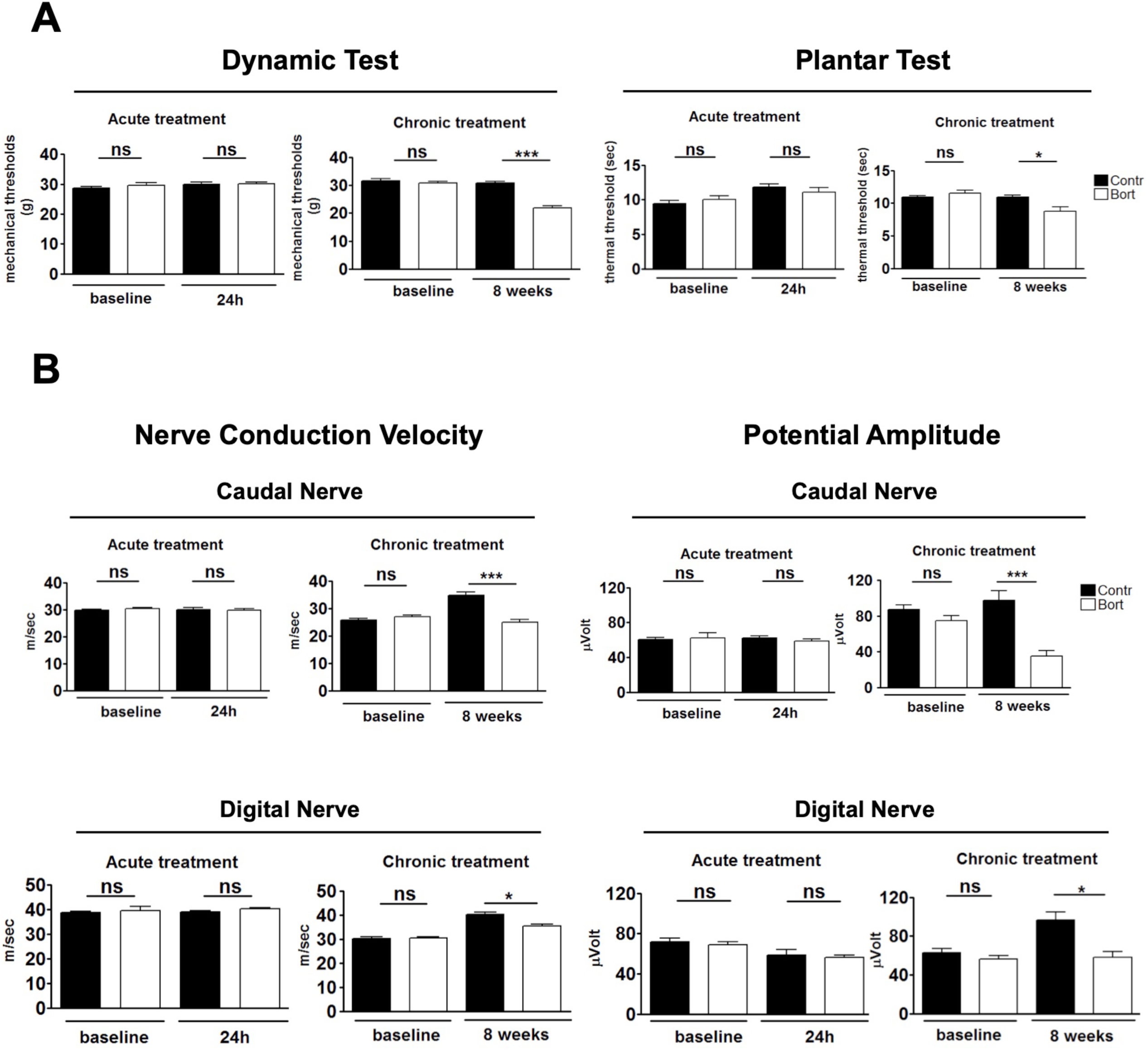
Behavioral tests and neurophysiological studies in control and Bort-treated rats after 24h or 8w of treatment (**A**) Hind-paw withdrawal response to mechanical stimulus and plantar withdrawal latency response to heat source were recorded before chemotherapy administration (baseline value), after 24h and at the end of treatment (8w) with Bortezomib (Bort). Mechanical allodynia and thermal hyperalgesia were observed only at the end of chronic administration of Bort in treated animals (Bort) compared to controls (Contr). (**B**) Digital and caudal nerve conduction velocity (NCV) and amplitude were measured before chemotherapy administration (baseline value), after 24h and at the end of treatment (8w) with Bort. Only after 8w of treatment with Bort significant decrease of caudal and digital NCV and potential amplitude were evident in treated rats vs controls (Contr). Data are expressed as mean values ±SEM, *** p<0.0001; * p<0.01 by t-test (unpaired, two-tailed); n=8 animals per condition.

**SFigure 2.**
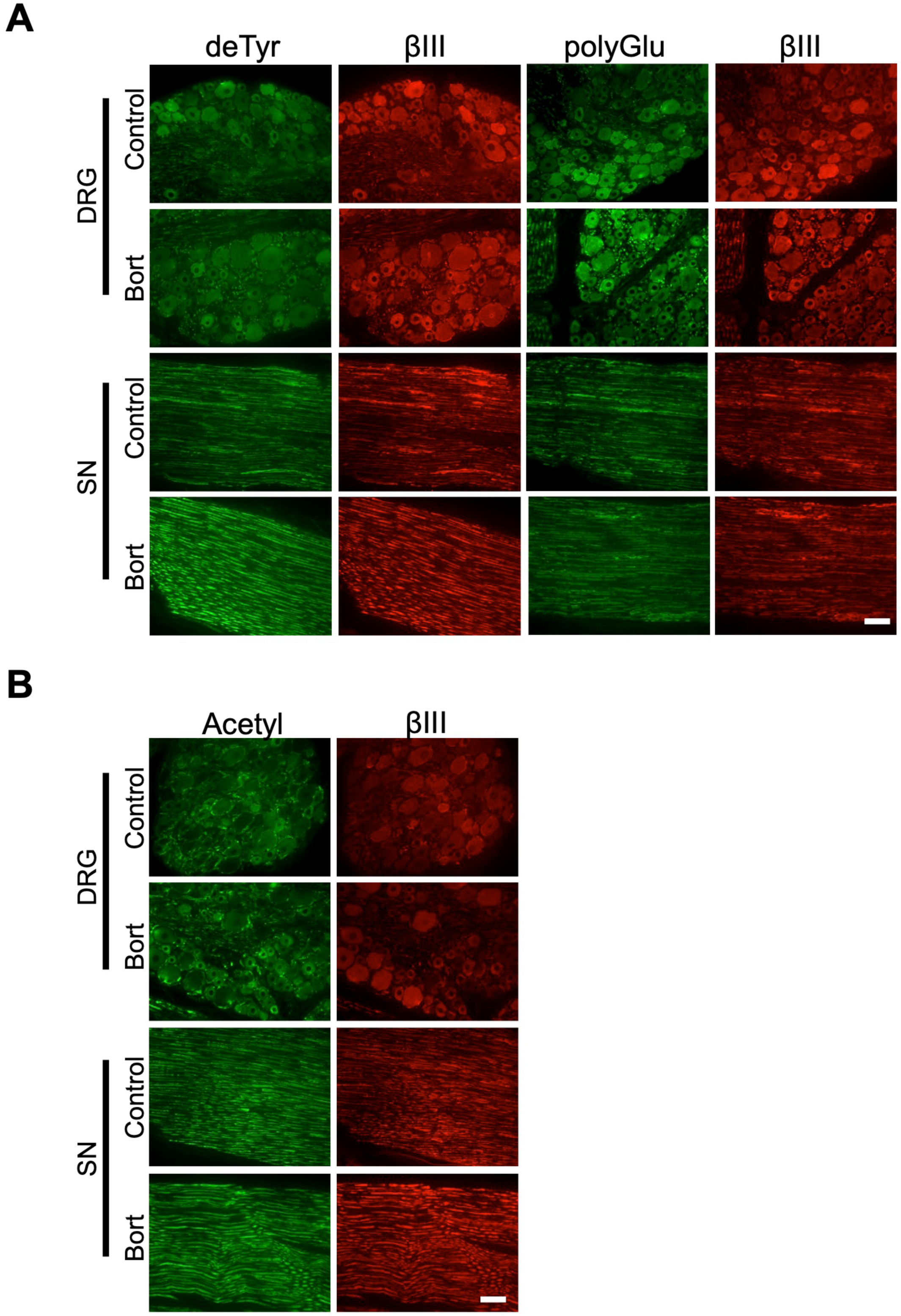
(A, B) Representative tubulin PTM immunofluorescence staining of DRG cell bodies and sciatic nerve (SN) isolated from rats acutely treated with Bortezomib (Bort). deTyr, detyrosinated tubulin; polyGlu (GT335), polyglutamylated tubulin, Acetyl, acetylated tubulin, βIII, betaIII tubulin isoform. Bar, 20μm.

**SFigure 3.**
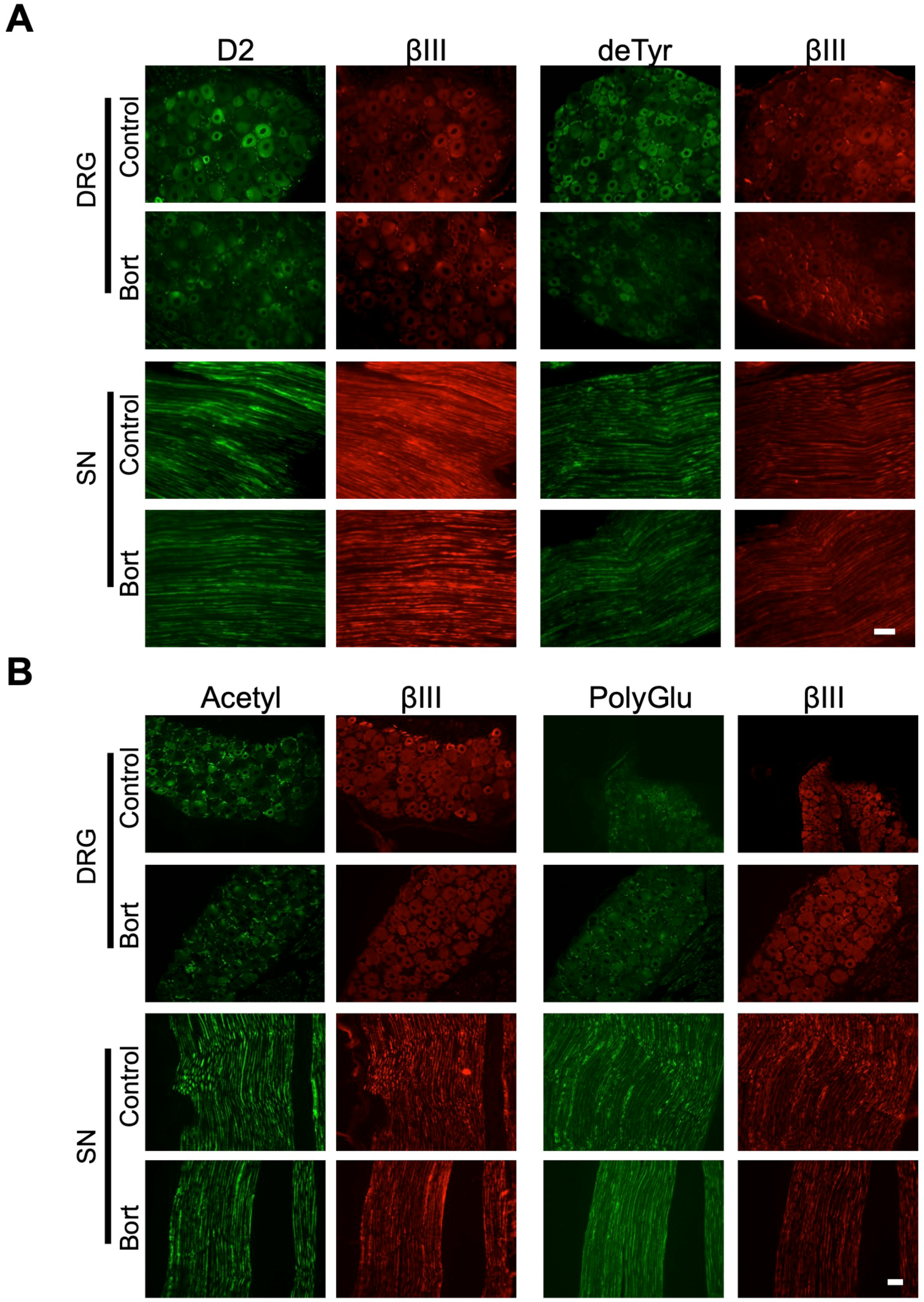
(A, B) Representative tubulin PTM immunofluorescence staining of DRG cell bodies and sciatic nerve (SN) isolated from rats chronically treated with Bortezomib (Bort). D2, delta 2 tubulin; deTyr, detyrosinated tubulin; polyGlu (GT335), polyglutamylated tubulin, Acetyl, acetylated tubulin, βIII, betaIII tubulin isoform. Bar, 50μm.

**SFigure 4.**
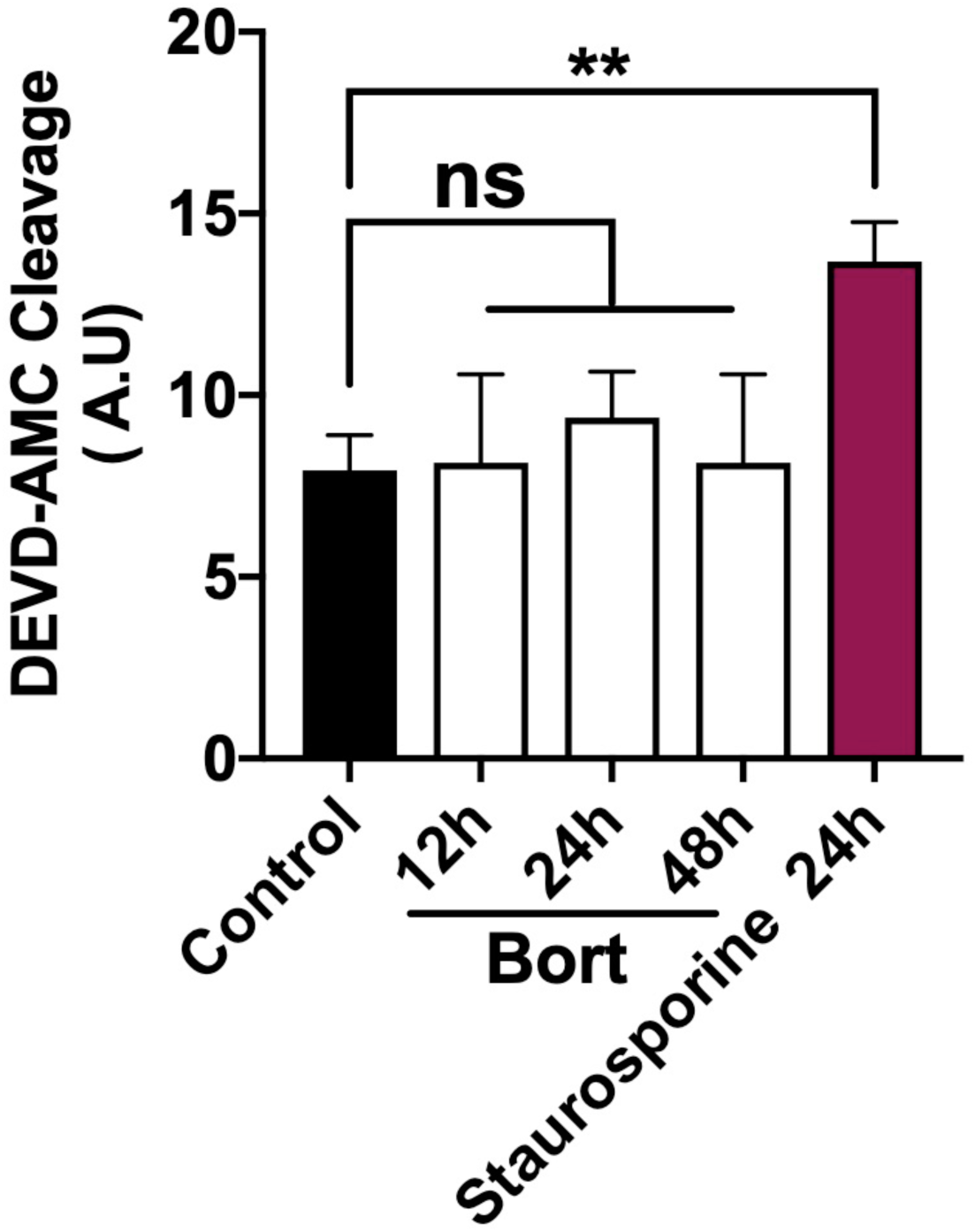
Caspase 3-like enzymatic activity was measured in DRG neurons treated with Bortezomib (Bort) at the concentration of 100nM. Staurosporine at the concentration of 600nM was added for 24h as a positive control. Data were reported as mean ± SEM (n=3). **p<0.001 by ANOVA.

**SFigure 5.**
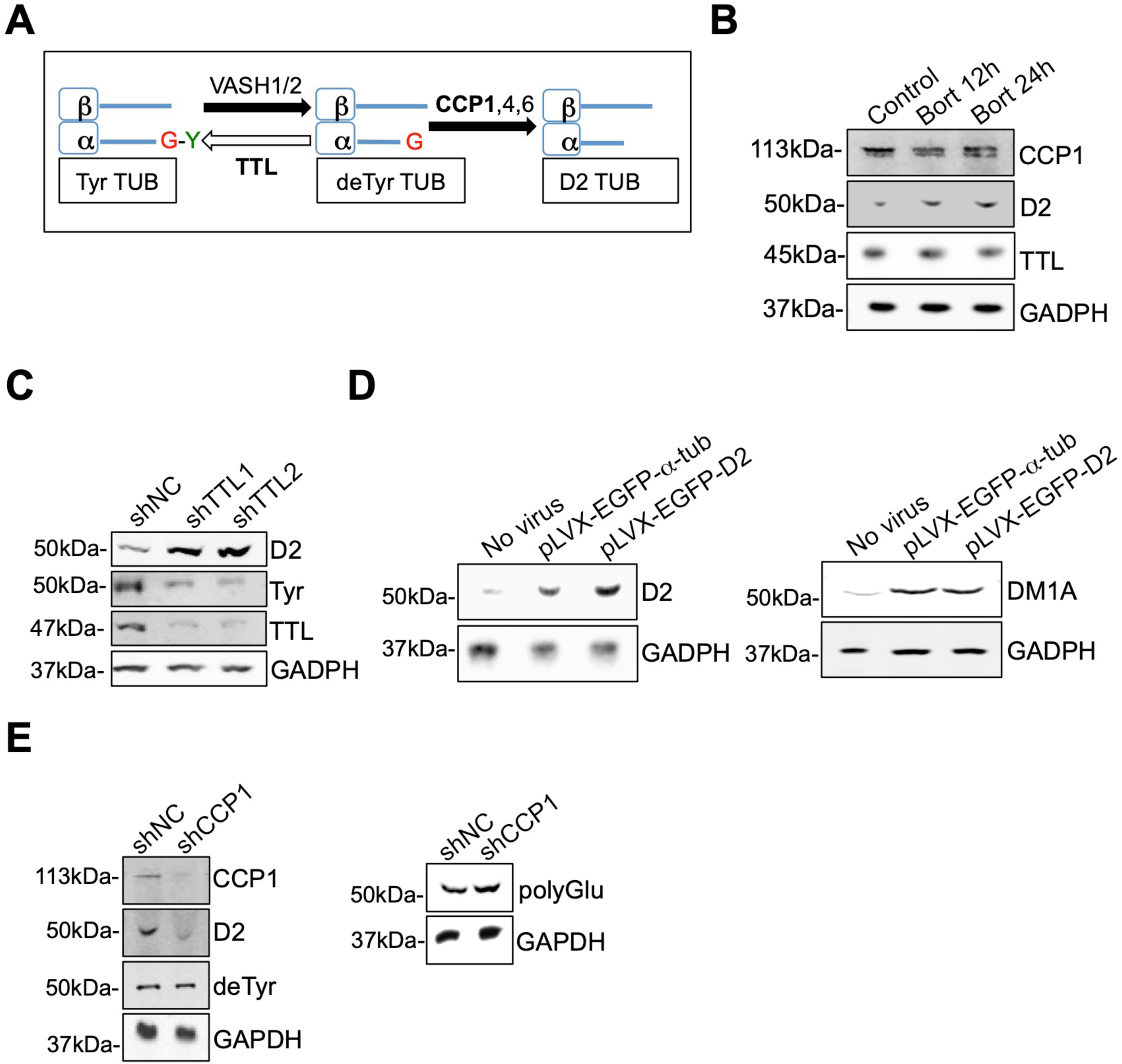
(A) Schematic of the enzymes involved in D2 generation. **(B)** IB analyses of carboxypeptidase 1 (CCP1), tubulin tyrosine ligase (TTL) and D2 levels in lysates from DRG neurons (12DIV) treated with 100nM Bort for the indicated times. **(C)** IB of tubulin tyrosine ligase (TTL), D2 and tyrosinated (TYR) tubulin levels in adult DRG neurons (12DIV) silenced of TTL expression by lentiviral shRNA delivery (5DIV). **(D)** IB of D2 and DM1A levels in adult DRG neurons (12DIV) expressing D2-(pLVX-EGFP-D2) or WT-tubulin (pLVX-EGFP-ααtub) (5DIV). **(E)** IB analyses of carboxypeptidase 1 (CCP1), polyglutamylated (polyGlu), D2 tubulin and detyrosinated tubulin (deTyr) levels in whole cell lysates from DRG neurons (5DIV) transduced with shRNA for CCP1 (1DIV). GAPDH, loading control.

**SFigure 6.**
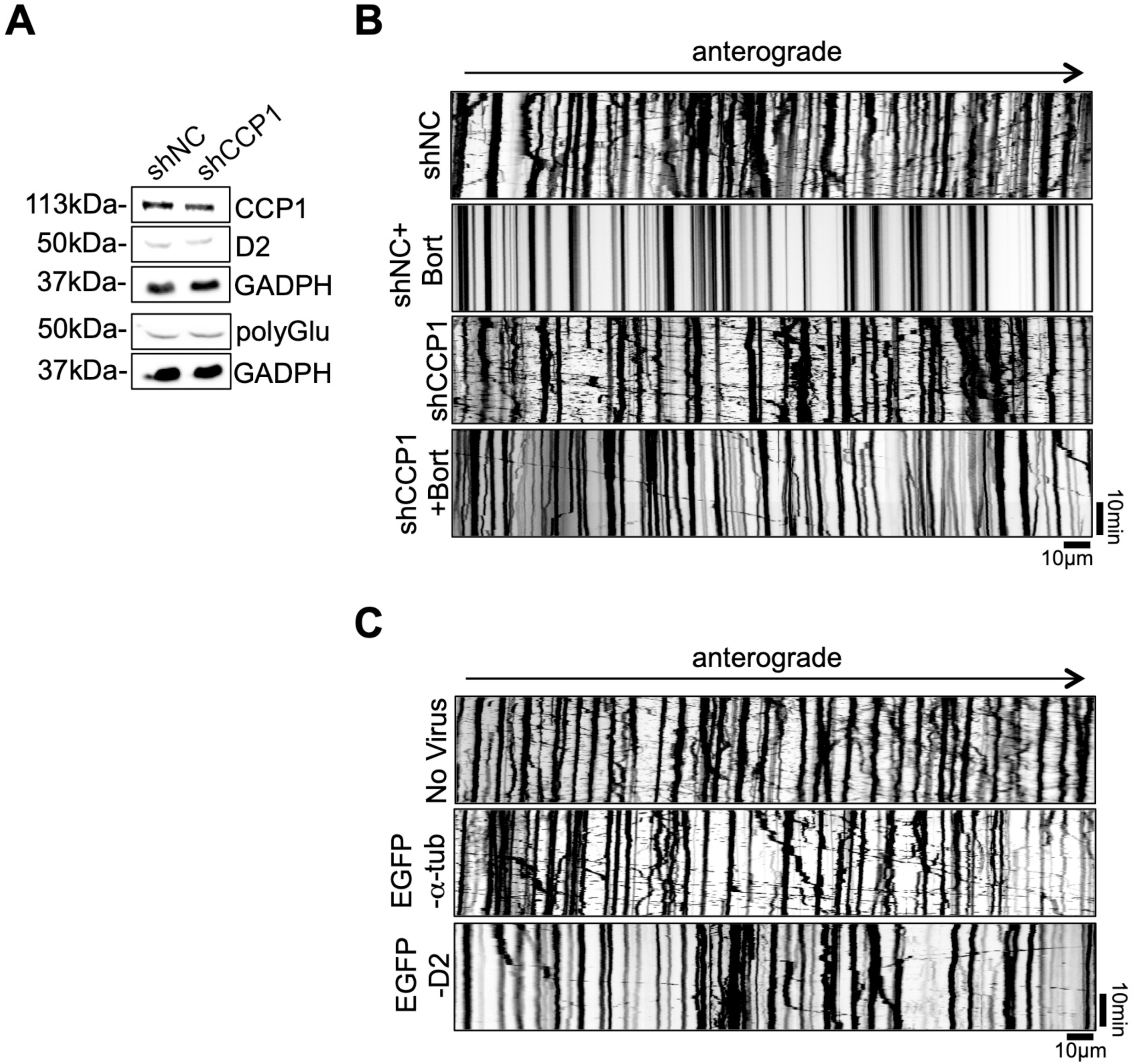
(A) IB of CCP1, D2, and polyglutamylated (polyGlu) tubulin levels in adult DRG neurons (5DIV) silenced of CCP1 expression at 1DIV for 5d. GAPDH, loading control. **(B)** Representative kymographs of mitochondria motility in DRG neurons (5DIV) infected at 1DIV with shCCP1 or shNC lentivirus prior to 72h infection with mito-DsRed lentivirus followed by Bort treatment (100nM for 24h). **(C)** Representative kymographs of mitochondria motility in DRG neurons (5DIV) infected at 1DIV with pLVX-EGFP-D2- or WT-tubulin lentivirus prior to 72h infection with mito-DsRed lentivirus. Videos (10s/frame for 30min).

## AUTHOR CONTRIBUTIONS

M.E.P., F.B. and G.C. have conceived the project. M.E.P. and X.Q. have carried out and analyzed all the *in vitro* experiments and analyzed the *in vivo* specimens. A.K. has contributed to the biochemical characterization of cultured DRG neurons and in the generation of molecular biology reagents. C.M. and L.M have carried out all the *in vivo* treatments and behavioral analysis. P.A. and G.F. have performed the neurophysiology experiments. M.S. and M. R. have performed and analyzed the experiments in zebrafish. K.T. and T.H.B. have provided human biopsies and clinical history details on the patients. M.E.P. and F.B. have written the manuscript.

## ACKNOWLEDGMENTS

This project was funded by the Thompson Family Foundation (TFFI) to F.B. and G.C. and by an RO1AG050658 (NIH/NIA) to F.B.. We are grateful to Gregg G. Gundersen for stimulating discussions and access to his microscopes. We thank Grace Shin for helpful discussions, Samie Jules for technical assistance in the isolation and analysis of the experiments with adult DRG neurons, and Annalisa Canta, Alessia Chiorazzi, Elisa Ballarini and Virginia Rodriguez for technical assistance in carrying out the *in vivo* experiments. The authors declare no competing financial interests.

